# Tuning intercellular cohesion with membrane-anchored oligonucleotides

**DOI:** 10.1101/620989

**Authors:** Ian T. Hoffecker, Yusuke Arima, Hiroo Iwata

## Abstract

Cohesive interactions between cells play an integral role in development, differentiation, and regeneration. Existing methods for controlling cell-cell cohesion by manipulating protein expression are constrained by biological interdependencies, e.g. coupling of cadherins to actomyosin force-feedback mechanisms. We use oligonucleotides conjugated to PEGylated lipid anchors (ssDNAPEGDPPE) to introduce artificial cell-cell cohesion that is largely decoupled from the internal cytoskeleton. We describe cell-cell doublets with a mechanical model based on isotropic, elastic deformation of spheres to estimate the cohesion at the cell-cell interface. Physical manipulation of cohesion by modulating PEG-lipid to ssDNAPEGDPPE ratio, and conversely treatment with actin-depolymerizing cytochalsin-D, resulted respectively in decreases and increases in doublet contact area. Our data are relevant to the ongoing discussion over mechanisms of tissue surface tension and in agreement with models based on opposing cortical and cohesive forces. PEG-lipid modulation of doublet geometries resulted in a well-defined curve indicating continuity, enabling prescriptive calibration for controlling doublet geometry. Our study demonstrates tuning of basic doublet cohesion, laying the foundation for more complex multicellular cohesion control independent of protein expression.

## Introduction

Cell-cell cohesion is increasingly recognized as a important practical control parameter in tissue engineering applications^1–3^ and in guidance of cell behaviors such as differentiation^4–6^. However cell-cell cohesion mediated by membrane proteins (cadherins, integrins, and the immunoglobulin superfamily) are inextricably connected to other biochemical processes. Cadherins interface with cytoskeletal actin on the cytosolic side of the membrane associating with force generating processes like migration, mechanosensing, and ECM adhesion^7–10^. Artificial control of protein-mediated cohesion can cause unintended consequences in coupled processes such as differentiation, proliferation, and apoptosis^11–13^ demanding that one negotiate such interdependency by fixing variables to reduce degrees of freedom^14–16^.

Manipulating expression of cohesion-mediating proteins remains the dominant strategy for modulating cohesive interactions between cells^17–19^, though a converse strategy is to restrict the spatial positioning of cells with 2D and 3D localization methods such as micropatterning to induce natural cohesion in a predictable manner^20–24^.

Cell surface modification with phospholipid-anchored oligonucleotides (ssDNAPEGDPPE) offers a means of inducing physical attachment of cells to substrates or other cells orthogonal to natural adhesion and cohesion biochemistry^25–28^. Recently, Y. Teramura showed that ssDNAPEGDPPE could be used to investigate the interactions of cohering cell doublets^29^ demonstrating dependence of cell-cell interactions on their degree of contact. We have also previously shown that native integrin-mediated adhesion modes take place in parallel with artificially-induced attachment with ssDNAPEGDPPE^28^.

The goal of this study is to use artificial ssDNAPEGDPPE-mediated cohesion in a cell doublet model as a tool to manipulate geometric parameters, e.g. their interfacial contact area and extent of membrane deformation, while minimizing the effects of coupling to mechanical, cytoskeletal biochemistry characteristic of other cohesion modes. We show that by tuning the concentration of ssDNAPEGDPPE molecules on the surface of cells before doublet formation, it is possible to achieve a well defined range of doublet geometry distributions characterized by interfacial contact area.

## 1 Materials and Methods

### 1.1 supplies and reagents

*α*-N-hydroxysuccinimidyl-*ω*-maleimidyl poly(ethylene glycol) (NHS-PEG-Mal, MW 5000), *α*-Succinimidyl carbonyl-*ω*-methoxy, polyoxyethylene (PEGDPPE), and 1,2-dipalmitoyl-sn-glycerol-3-phosphatidylethanolamine (DPPE) were purchased from NOF Corporation (Tokyo, JP). Fetal bovine serum (FBS), Hanks Balanced Salt Solution (HBSS), and RPMI 1640 medium were purchased from Invitrogen, Co. (Carlsbad, CA, USA). Cytochalsin D was purchased from Sigma Aldrich(St. Louis, MO, USA).

Single stranded DNA (ssDNA) with 5’ terminal disulfide modifications and ssDNA with 5’ disulfide and 3’ fluorescein amidite were purchased from Fasmac (Atsugi City, Kanagawa Japan).

**Table 1.**
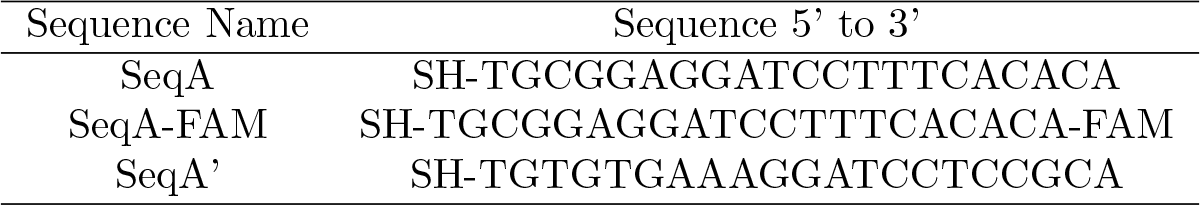
Thiol-modified ssDNA Sequences

### 1.2 modification of cells with ssDNAPEGDPPE

The cells used in this experiment were CCRF-CEM cells, a human T-cell lymphoblast-like cell line and non-adherent cell type used in previous ssDNAPEGDPPE experiments^30^. Oligo-terminated phospholipids (ssDNAPEGDPPE) (Fig1 a)^30,31^ were synthesized in two steps as previously described by first conjugating NHS-PEG-maleimide to DPPE followed by a second conjugation of maleimide-PEG-DPPE to thiol-terminated oligonucleotides. PEGDPPE was produced with an analogous single step conjugation of PEGNHS to DPPE. Cells were modified with ssDNAPEGDPPE by incubation for 1 *hr*, RT, in a solution of ssDNAPEGDPPE at a concentration of 200 *μg/mL*. Cells were rinsed thrice with HBSS followed by pelleting. Doublets were formed by mixing complementary ssDNAPEGDPPE-bearing cell groups into a single glass bottom dish during observation with confocal microscopy. Oligonucleotides anchored to cells were visualized by substituting SeqA-PEG-DPPE for fluorescein-terminated FAM-SeqA-PEG-DPPE.

**Figure 1.**
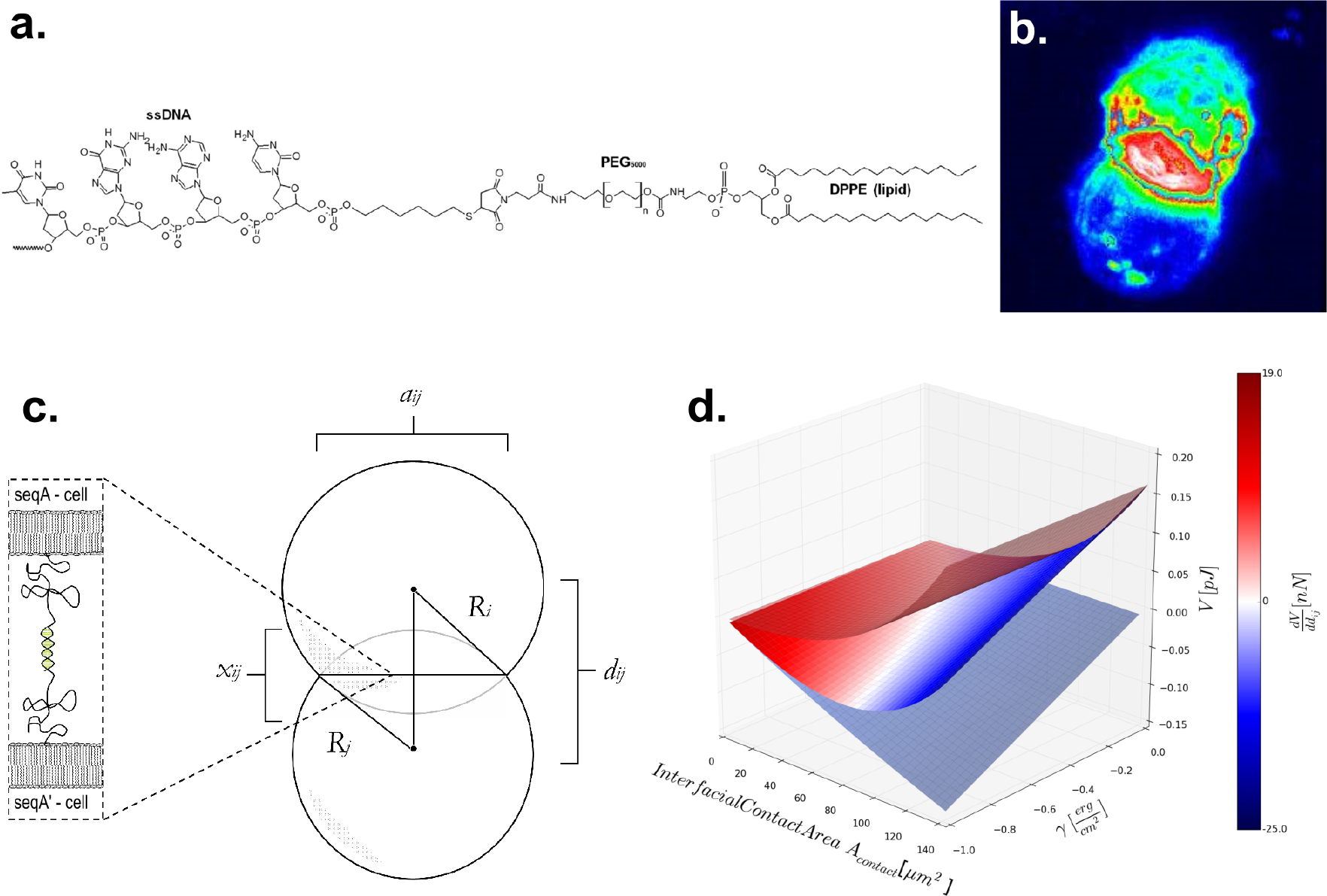
(a) Chemical structure of ssDNAPEGDPPE. (b) Tomographic reconstruction of fluorescently labeled cell doublet cohering through FAM-modified ssDNAPEGDPPE hybridization across adjacent membranes. (c) Coarse-grained geometric representation of an ideal cell doublet: 2 intersecting spheres with truncated regions corresponding to deformed elastic cells. (d) Surface plot of potential energy as a function of doublet interfacial contact area and interfacial cohesion and color map representative of the force along the center-to-center axis.

### 1.3 formation of cell doublets

Cell doublets were formed after preparing two parallel populations of modified cells bearing respectively (FAM)-SeqAPEGDPPE and its complement SeqA’PEGDPPE such that modified cells could form complementary pairings upon mixing of the two populations. Complementary cell groups were exposed to each other over a glass bottom dish during imaging by confocal laser scanning microscopy and allowed to settle after aggregation to reach steady state over the course of 15 *min*. Doublets were identified manually out of a population also containing single cells, triplets, and larger aggregates. A tomographically reconstructed 3D image of a ssDNAPEGDPPE-cohered cell doublet with fluorescently labeled ssDNA is shown in Fig 1 b.

For cells modified to exhibit reduced cohesion, PEGDPPE was added to the ssDNAPEGDPPE solutions during the cell modification stage in various molar ratios to provide binding competition and reduce the equilibrium ssDNAPEGDPPE membrane surface concentration. Co-PEGDPPE/ssDNAPEGDPPE-modified cells were combined in complementary pairs as with ssDNAPEGDPPE-modified cells to form a distribution composed of single cells, doublets, triplets, and high order cell-cell aggregates from which doublets were manually identified and imaged with confocal microscopy.

For cells treated pharmacologically to exhibit reduced elastic repulsion, cytochalasin-D (1 *mg/mL* in DMSO) was added to cell suspensions during the 1 *hr* ssDNAPEGDPPE modification for a final concentration of 2 *μg/mL*^32^, and cells observed with confocal microscopy were kept in medium containing 2 *μg/mL* cytochalasin-D.

### 1.4 estimation of cohesion via doublet geometry

Cells *i* and *j* interact with a total energy

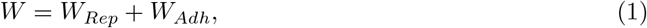

where *W*_*Rep*_ is the potential energy contribution due to elastic repulsion and *W*_*Adh*_ is the potential energy contribution resulting from adhesion over the surface of contact.

To estimate the adhesion energy, we have chosen to model the cell repulsive energy due to elastic deformation by the Hertz-model applied to two compressed elastic spheres described in Landau and Lifschitz’s Theory of Elasticity^33,34^ shown schematically in Fig. 1 c. Two cells *i* and *j* of unequal spherical radii *R*_*i*_ and *R*_*j*_ are divided by an interfacial contact area *A*_*contact*_ of diameter *a*_*ij*_. The two cell centers of curvature are separated by a distance *d*_*ij*_, and the truncated caps due to adhesion are together of length *x*_*ij*_ referred to as the indentation depth. The contribution to interfacial energy due to repulsion is

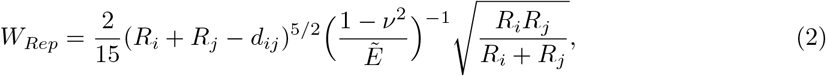

where 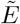 and *ν* are the Young’s modulus and Poisson’s ratio respectively defined for both cells *i* and *j*.

We assume that the radii of curvature of cells can be represented by spherical radii. We imported the Young’s modulus, which we assume for simplicity to be constant in the Hertz model, from Rosenbluth et al. (*E* = 855 ±670 *Pa* for HL60 leukemia myeloid cells, and likewise assumed a Poisson’s ratio of 0.5 as was done so by Rosenbluth et al^35^.

The adhesive contribution to potential energy is

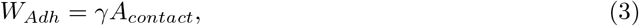

where *γ* is the cohesion represented in units of energy per unit area. The contact area can be rewritten as a function of *R*_*i*_, *R*_*j*_, and *d*_*ij*_ such that the potential due to adhesion becomes

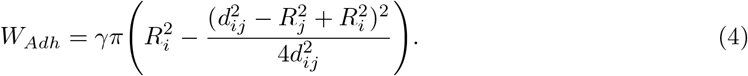

Thus, given values of *R*_*i*_, *R*_*j*_, and *γ*, the total interfacial energy between cells *i* and *j* can be determined as a function of *d*_*ij*_ or *x*_*ij*_. Fig. 1 d shows a plot of potential energies *W* (magenta), *W*_*Rep*_ (red), and *W*_*Adh*_ (blue) as a function of interfacial contact area *A*_*contact*_ and cohesion *γ* for a hypothetical pair of fixed volumes *V*_*i*_ = *V*_*j*_ = 900 *μm*^3^.

The minimum of *W* can be determined by solving for the root of its derivative with respect to separation distance, the point where the force along the axis extending from cell center to center is zero:

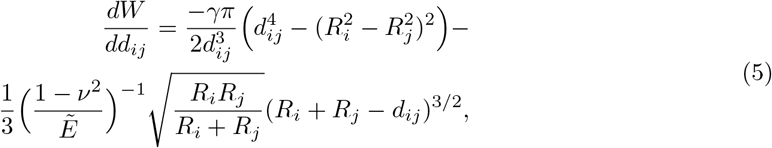

and is shown in Fig. 1 d as the color map. For values of *γ* sufficient to overcome thermal and convective noise, a potential energy well exists for *W* such that *A*_*contact*_ > 0 and cell cohesion occurs at steady state. This region is indicated by white on the color map in Fig. 1 d. The roots of the derivative equation can be rearranged to solve for cohesion:

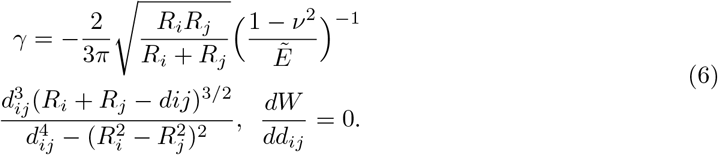

To compute the space of possible steady state doublet conformations including contact area as a function of both Young’s modulus and cohesion (Fig 2 d), a numerical algorithm was implemented to identify the values of *A*_*contact*_ which satisfy the root of 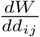 under an additional constraint of fixed cell volume (See Supplementary Methods).

**Figure 2.**
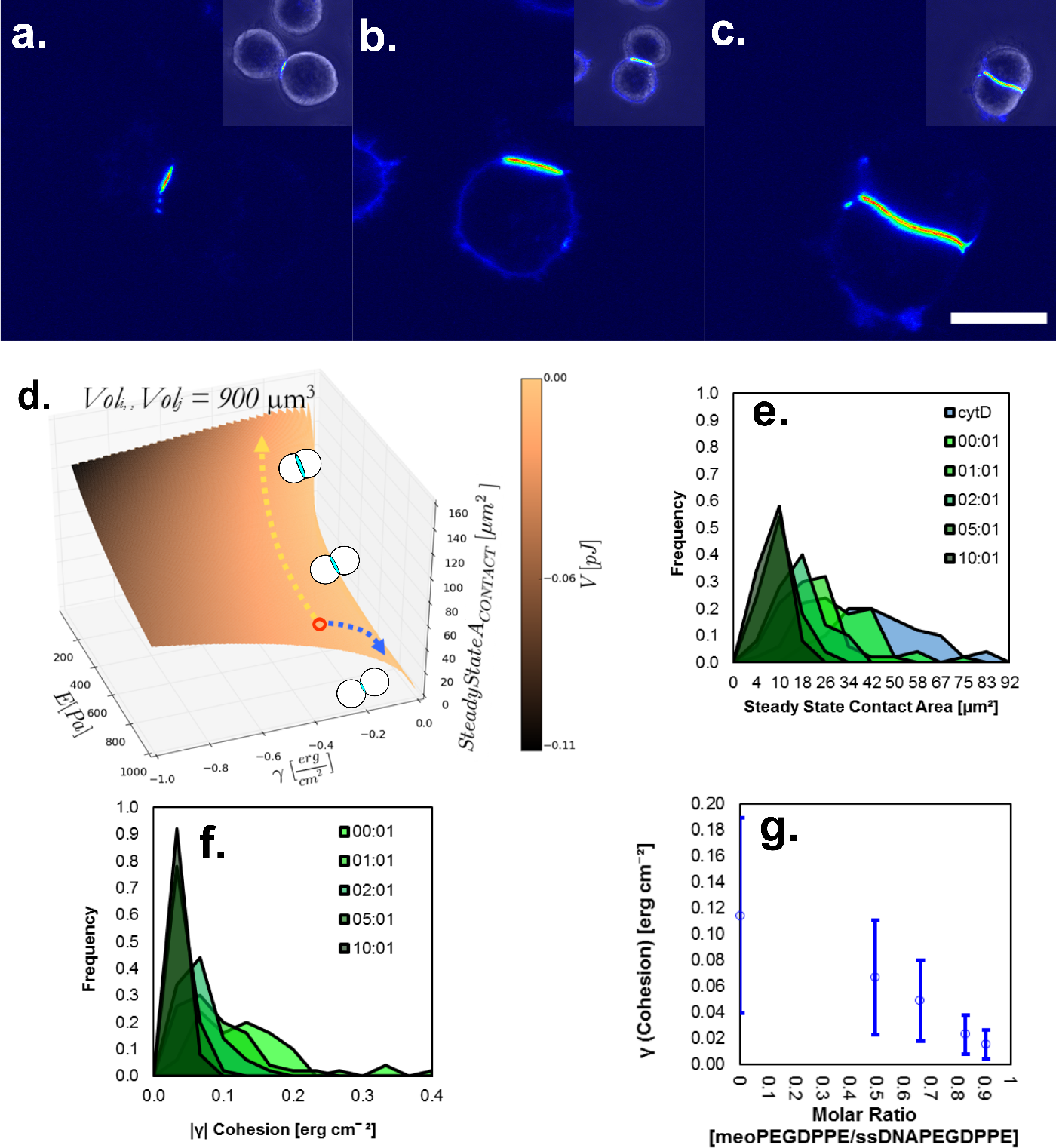
Area of the interfacial contact plane between cell doublets (*A*_*contact*_) can be influenced by modulating parameters: Young’s modulus or cohesion. (a) Cross-section of doublet formed from cells modified with a mixture of PEGDPPE and ssDNAPEGDPPE resulting in a greatly reduced *Ā*_*contact*_. (b) Normal doublet formed form cells modified with only DNA-PEG-DPPE. (c) Doublet with large *Ā*_*contact*_ formed from DNA-PEG-DPPE-modified cells treated with cytochalsin-D (d) Surface plot of *Ā*_*contact*_ (vertical axis) as a function of the Young’s Modulus 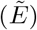 and the cohesion (*γ*) with the surface energy *W* indicated by the color map. Arrows indicate hypothetical changes in *Ā*_*contact*_ reached by orthogonal movements along the parameter axes. (e) Histogram of *Ā*_*contact*_ measurements determined from cross-sectional images of doublets formed under normal conditions, with the addition of PEGDPPE at various relative concentrations during cell modification, and with the addition of cytochalsin-D (blue). (f) Histogram of cohesion values calculated from the geometry of doublet cross sections based on a fixed and constant 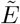 for doublets formed under normal conditions and doublets with various relative concentrations of PEGDPPE. (g) Plot of the absolute value of cohesion *|γ|* calculated on the basis of doublet geometry vs the molar fraction of PEGDPPE to ssDNAPEGDPPE used during cell modifications.

### 1.5 doublet imaging and processing

Mixed complementary groups of FAMSeqAPEGDPPE-modified and SeqA’PEGDPPE-modified cells were briefly agitated to induce aggregation by hybridization and pipetted onto glass bottom dishes and allowed to settle and reach a steady state for 15 minutes prior to imaging with a Fluoview FV500 confocal laser scanning microscope. Steady state was reached on the microscope sample stage to prevent further contact between unhybridized cells and cell aggregates. Doublets were identified manually and imaged with a 60x 1.35 NA oil objective. The z-depth of each cross-sectional image was adjusted manually to obtain an image of the maximally long central chord of the doublet interfacial contact area. While this procedure ensures unbiased measurement of the interfacial contact diameter *a*_*ij*_, it does not account for slanting of the axis between the two cells that could be caused by a difference in cell radii. An adjustment based on geometric constraints is applied then to values of *d*_*ij*_, *R*_*i*_, and *R*_*j*_ obtained experimentally (See Supplementary Data).

A partially automated image processing algorithm was applied to doublet images in order to extract their geometric parameters and the fluorescence intensity due to FAMSeqAPEGDPPE at interfaces (SFig 2). Cropped interface images (Fig 4 a) were then processed with a numerical signal detection algorithm (Supplementary Methods) which begins by shifting the alignment of rows of pixels along the vertical axis of the image such that the brightest pixel in each row is centered (Fig 4 b). This action corrects for irregularities in shape of the interface leading to a sharper distribution of pixel intensities when the image is collapsed into a single horizontal axis. The collapsed intensity profile peak is then integrated and divided by the interface diameter or chord length *a*_*ij*_ to obtain the intensity density of the profile in arbitrary intensity units.

**Figure 3.**
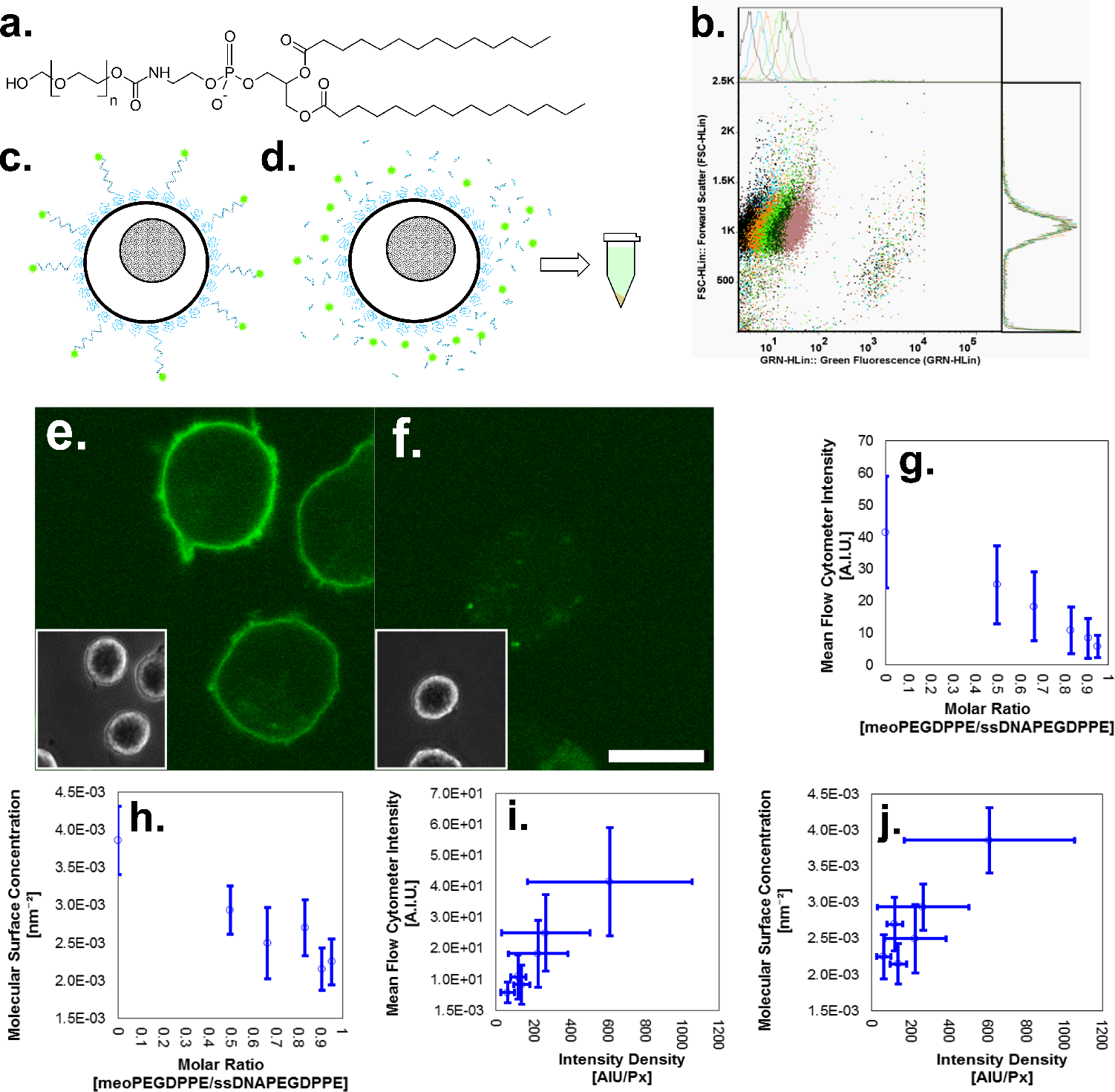
Modulation of PEGDPPE to ssDNAPEGDPPE ratio can be used to tune the density of hybridizable molecules on cell surfaces. (a) Structure of PEGDPPE. (b) Flow cytometry results displayed as forward scatter values vs green fluorescence. (c) Schematic illustration of a cell saturated with PEG-lipids, some of which terminate as methoxy-PEG while others are connected to FAM-labeled ssDNA. (d) Schematic illustration of FAM extraction by digesting nucleotides with Benzonase followed by cell separation from FAM-containing supernatant by centrifugation. (e) Before image of FAM-SeqA-PEG-DPPE-modified cells. (f) After image of FAM-SeqA-PEG-DPPE-modified cells following digestion with Benzonase. Scale bar = 10 *μm* (g) top: Example of a cell image cross-section transformed into a cartesian, axis-aligned profile for analysis of fluorescence intensity on the cell surface. bottom: Collapsed profile with peak corresponding to the total image fluorescence intensity signal along the cell periphery. (h) Plot mean of flow cytometry fluorescence measurements of cell populations modified with different relative molar fractions of PEGDPPE to FAMSeqAPEGDPPE. (i) Plot of population mean molecular surface concentrations computed on the basis of fluorescence spectroscopy of supernatants vs relative molar fraction of PEGDPPE to ssDNAPEGDPPE. (j) Plot population mean of flow cytometry fluorescence measurements vs population mean fluorescence intensity densities computed from confocal image analysis. (k) Plot of population mean molecular surface concentrations computed on the basis of fluorescence spectroscopy of supernatants vs population mean fluorescence intensity densities computed from confocal image analysis.

**Figure 4.**
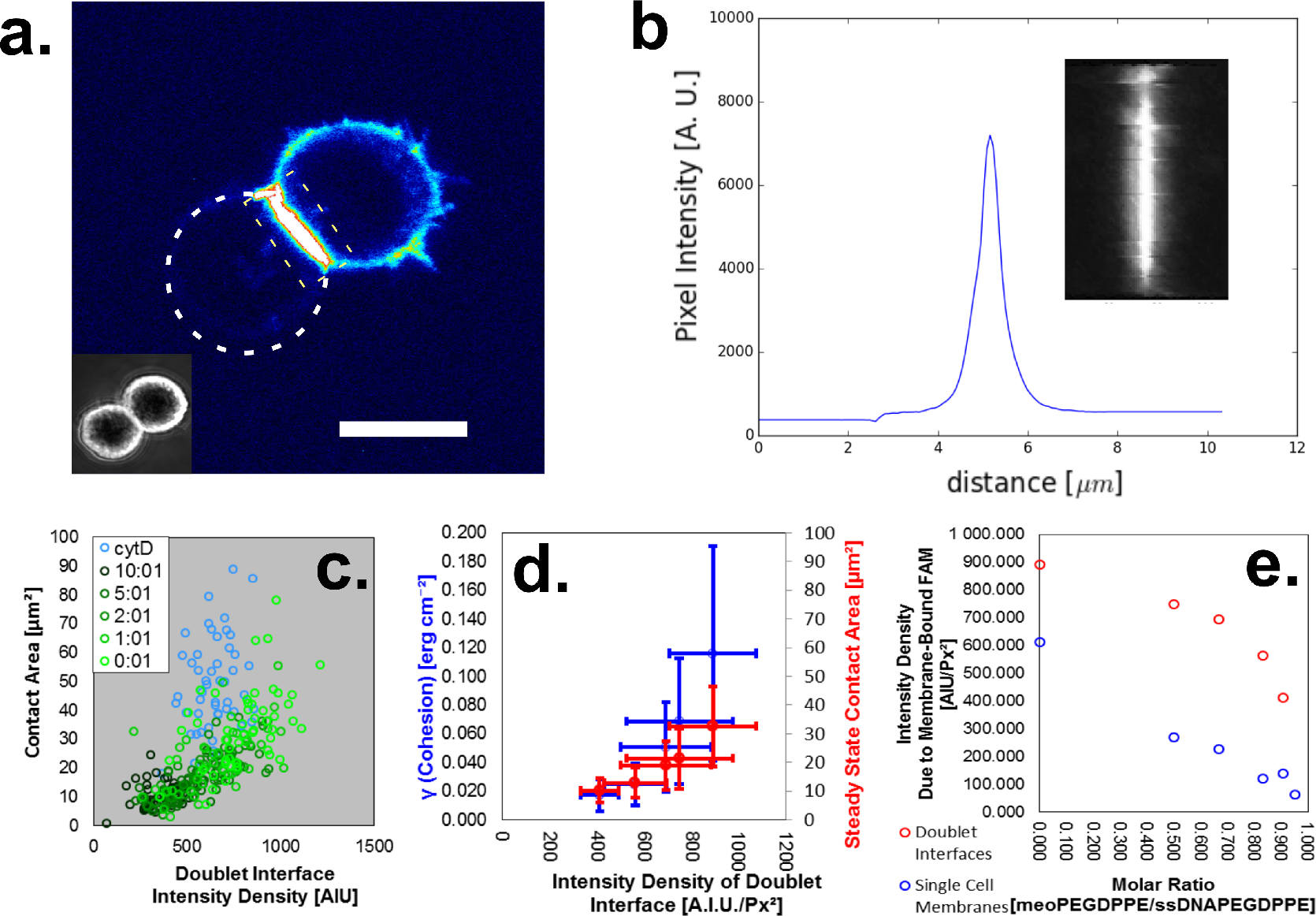
Image analysis of doublet interface cross-sections. (a) Raw image of a doublet interface cross-section taken with confocal microscopy. Scale bar: 10 *μm* (b) Doublet interface image from a. after pixel alignment. (c) Summed pixel intensities from b collapsed onto a single x axis. (d) Contact area measurements for cytochalsin-D treated doublets plotted against their computed intensity density values at different PEGDPPE:ssDNAPEGDPPE molar ratios. (e) Absolute value of cohesion population means and steady state contact area population means plotted versus their corresponding mean intensity densities. (f) Intensity density population means of image-processed doublet interfaces (red) and single cell membranes (blue) plotted as a function of the molar ratio of PEGDPPE to ssDNAPEGDPPE.

### 1.6 fluorescence spectroscopy of digested FAM-ssDNAPEGDPPE

UV/Vis spectroscopy was carried out using a Beckman DU-640 spectrophotometer (Beckman Coulter, Brea CA, USA) and fluorescence spectroscopy with a Hitachi F-2710 fluorometer (Hitachi, Tokyo, Japan). To calibrate the concentration estimates of prepared solutions of FAM-ssDNAPEGDPPE, a standard was constructed with UV/vis absorbance measurements taken at 260 *nm* and adjusted to account for the absorbance contribution from FAM. A linear calibration curve (SFig 3 a) was used to adjust concentration estimates used in later steps. A standard was then constructed with fluorescence spectroscopic measurements at several different concentrations of FAMSeqAPEGDPPE digested with Benzonase (1x RQ1 buffer (Promega, Madison, WI, USA) supplemented 0.9% NaCl and 2500 *U/mL* Benzonase, and incubated for 10 minutes at 37°C) to mimic the conditions of cell experiments requiring the liberation of FAM from FAMSeqAPEGDPPE-modified cell membranes (SFig 3 b). The coefficients of the linear standard, *k*_*spec*_ = 3.71 ± 0.04 · 10^−4^ for the slope and *β*_*spec*_ = 9.59 ± 1. · 10^−2^ for the intercept, were then used to infer the FAM concentrations of supernatants extracted from suspensions of cells bearing FAMSeqAPEGDPPE following DNA digestion with Benzonase.

Suspensions of FAMSeqAPEGDPPE-bearing cells were first prepared at concentrations of approximately 200 *cells/μ*, washed, pelleted, and resuspended in 1x RQ1 buffer (Promega, Madison, WI, USA) supplemented 0.9% NaCl and 2500 *U/mL* Benzonase and incubated for 10 minutes at 37°C. The cells were again pelleted and the supernatant containing liberated FAM and digested nucleotide fragments was extracted for spectroscopic analysis. Prior to digestion, a fraction of suspensions were analyzed with flow cytometry to estimate the total number of cells in the Benzonase treated sample. Supernatant fluorescence spectroscopy was performed using 495 *nm* excitation with a detection range spanning 510 to 550 *nm*.

### 1.7 flow cytometry analysis of FAM-ssDNAPEGDPPE-modified cells

Cells sampled from the same non-aggregated populations of FAMSeqAPEGDPPE-modified cells as those analyzed by confocal microscopy and post-digestion fluorescence spectroscopy were sampled with a flow cytometer (5000 cells, Guava Easy Cyte Mini, Millipore, Billerica, MA, USA). Flow cytometry-based measurement of number of cells per *μL* of suspension were used to determine the number of cells treated with Benzonase for fluorescence spectroscopy analysis of liberated FAM.

### 1.8 single cell imaging and membrane fluorescence profile quantification

Confocal microscopic profiles of non-aggregated single cells modified with FAMSeqAPEGDPPE were analyzed in a semi-automated fashion. Manually identified centers of the cells in cross-section images were used as the origin of a polar to Cartesian transformation of the fluorescent halos visible along cell membranes with intensities indicating relative amounts of FAMSeqAPEGDPPE on the cell surface. Fluorescence profiles were aligned on a single vertical axis to increase the sharpness of profile peaks prior to integration of the peak for single quantification of the fluorescence intensities (See Supplemental Materials and Methods).

### 1.9 Statistics

Error propagation according to the variance equation was estimated for molecular densities, cohesion measured via geometry, and steady state interfacial contact area values. Error bars for these computed quantities represent propagated errors, and error bars in all other cases indicate standard deviations.

## 2 Results

### 2.0.1 development of coarse-grained cell doublet model

By modifying complementary batches of cells respectively with oligonucleotide-conjugated PEG-lipids (SeqAPEGDPPE and SeqA’PEGDPPE; Fig 1 a) followed by suspending the two populations together, we formed doublets along with other aggregates of CCRF-CEM cells cohered at an adjoining interface by the hybridization of complementary single stranded oligonucleotides anchored to the cell membranes. By substituting SeqAPEGDPPE with a fluorescently labeled analog (FAMSeqAPEGDPPE), it was possible to visualize the distribution of ssDNAPEGDPPE of one of the cells in each doublet and construct a tomographic picture from multiple confocal laser scanning microscopic sections (Fig 1 b). Based on this information and the precedent set by similar models of cadherin-mediated cohesion^36–39^, we have developed a coarse-grained model of the DNAPEGDPPE-cohered cell doublet consisting of two intersecting spheres of unequal radii and plane of intersection representing the interface of cohesion between two deformed elastic spheres (Fig 1 c). The dynamics of the doublet is driven by the imbalance of repulsive elastic force and attractive cohesion force in the direction of the center-to-center axis (Equation 5), and when interfacial cohesion is greater than zero, a non-zero interfacial contact area exists corresponding with the potential energy minimum where repulsive and attractive forces are balanced (Fig 1 d).

We designed and implemented a numerical algorithm to compute the interfacial contact area *A*_*contact*_ at steady state (*Ā*_*contact*_), a key geometric parameter indicative of the extent of cohesive deformation for a given cell type of known size distribution. The interfacial contact area *A*_*contact*_ is at steady state when attractive and repulsive forces are balanced for two cells of fixed volume cohered according to our doublet model as a function of the Young’s modulus 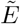 and the cohesion parameter *γ* or adhesion energy per unit area at the interface. This dependence is shown as a surface plot in Fig 2 d. Examining the space of *Ā*_*contact*_, we can see that doublets formed with low *γ* and large 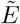 are smallest corresponding to weakly adhesive stiff cells. Conversely, doublets with large *γ* and low 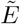, in other words soft and highly adhesive cells, have the largest predicted *Ā*_*contact*_ and deepest potential energy well indicated by black on the color map in Fig 2 d. To test our model, we wished to explore this space of *Ā*_*contact*_’s by modulating *γ* and 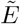 independently of each other.

### 2.0.2 experimental investigation of independent elastic and cohesive parameters of doublet geometry

We prepared cell doublets with reduced interfacial cohesion by reducing the surface concentration of ssDNAPEGDPPE through the introduction of non-hybridizing methoxy-PEG-lipids (PEGDPPE; Fig 3 a) during the cell-modification stage to compete with ssDNAPEGDPPE for binding sites on the cell membrane (Fig 3 c). When cells were modified in buffer containing a 10:1 molar ratio of PEGDPPE to FAMSeqAPEGDPPE/SeqA’PEGDPPE (Fig 2 a), confocal images of resulting doublets revealed cross-sections of *Ā*_*contact*_ values qualitatively smaller than those of normal doublets prepared from cell groups modified without PEGDPPE (Fig 2 b).

We wished to reduce the mechanical integrity of cells, so we treated cells during the ssDNAPEGDPPE modification stage and during doublet formation/imaging with cytochalsin D, a cytoskeletal inhibitor previously shown prevent actin polymerization and reduce cortical integrity^40,41^. Doublets prepared with cytochalsin D (Fig 2 c) exhibited qualitatively larger *Ā*_*contact*_ cross-sections in confocal images compared to normal doublets (Fig 2 b).

In addition to the 10:1 PEGDPPE:ssDNAPEGDPPE, normal (0:1 PEGDPPE:ssDNAPEGDPPE), and cytochalsin D-treated preparations, we prepared doublets from cells modified with 5:1, 2:1, and 1:1 PEGDPPE:ssDNAPEGDPPE molar ratios and analyzed confocal images of n=50 doublets per condition by manually tracing circles along the contours of doublet cross-sections from which the geometric parameters *d*_*ij*_, *R*_*i*_, and *R*_*j*_ could be computed. From these values, interfacial diameters *a*_*ij*_ and *Ā*_*contact*_ of each doublet were computed and compiled into the histogram shown in Fig 2 e. Provided our assumed values for the Young’s modulus and Poisson’s ratio, we computed the cohesion from the assumption that cohesive and elastic forces are balanced (Equation 6) and compiled these into a histogram in Fig 2 f. Since we do not know 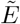 of the cytochalasin-D-treated doublets relative to our postulated 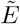, this condition was excluded from cohesion computation. Despite large standard deviations in ensemble cohesion estimates (SE > 60%), strong correlation between the molar ratio of PEGDPPE:ssDNAPEGDPPE and the cohesion computed on the basis of our geometric model and large sample sizes permits prescriptive control over the average doublet geometry (SEM < 10%) (Fig 2 g).

### 2.1 fluorescence-based quantification of hybridizable molecular surface concentration

In our mechanical model of the cell doublet, we infer the cohesion at the cell-cell interface on the basis of force balance with elastic forces in absence of knowledge of the molecular distribution. To better understand the distribution of ssDNAPEGDPPE molecules at the cell-cell interface and their contribution to, we utilize the fluorescence of fluorescein-terminated FAMSeqAPEGDPPE to track the distribution and concentration of ssDNAPEGDPPE molecules in the cell membrane. We first examined single cells prepared with mixtures of PEGDPPE and FAMSeqAPEGDPPE with flow cytometry (Fig 3 b). Forward scatter vs green fluorescence distributions were of highly reproducible shape between conditions besides being shifted in their peak fluorescence according to the PEGDPPE:FAMSeqAPEGDPPE ratio used in their preparation. The molar ratio of PEGDPPE to SeqAPEGDPPE correlates inversely with the mean peak fluorescence of cell populations analyzed with flow cytometry (Fig 3 g) demonstrating prescriptive control over the relative mean concentration of ssDNAPEGDPPE on cell surfaces through the addition of PEGDPPE during modification.

We developed a fluorescence spectroscopy-based assay to estimate the surface concentration of FAMSeqAPEGDPPE on membranes of single cells sampled from the same population of cells used for flow cytometry. We used oligonucleotides terminated with a single fluorescein moiety to ensure that the fluorescent emission corresponds to oligonucleotide (SeqA) concentration while leaving PEGDPPE molecules unlabeled (Fig 3 c). We used Benzonase, a nonspecific nuclease capable of digesting ssDNA and dsDNA, to digest away the oligonucleotide-tethered fluorescein molecules, releasing them into the surrounding medium which we collected and analyzed with fluorescence spectroscopy (Fig 3 d). Fluorescent halos marking the contour of the cell membrane in confocal cross-sections could be observed after cells were modified with FAMSeqAPEGDPPE (Fig 3 e), while after Benzonase digestion and removal of the digest supernatant the remaining cells exhibited negligible fluorescence levels when re-imaged (Fig 3 f) indicating that nearly all of the fluorescein was successfully extracted. Based on a linear calibration relating solution fluorescence intensity to the solution ssDNA concentration (see Supplementary methods) we measured the total amount of fluorescein extracted and could then estimate the average FAMSeqAPEGDPPE concentration per cell by dividing by the number of cells used in the extract preparation. As with flow cytometry-based fluorescence intensity, the mean molecular surface concentration correlates inversely with the PEGDPPE:FAMSeqAPEGDPPE molar ratio (Fig 3 h).

Sampling from the same populations of cells used for flow cytometry and fluorescence spectroscopy analyses, we also systematically gathered confocal images of cell cross-sections. Images were processed semi-automatically by first manually identifying the centers of each cell prior to a polar-Cartesian transform and an automated pixel alignment of the resulting linear membrane image. The vertically aligned membrane images were each collapsed into 1D intensity profiles that could be integrated for each cell analyzed (SFig 4). The polar-Cartesian transformations were non-conservative in that the density of pixel brightness rather than total pixel values were preserved, thus integrated membrane profile values correspond to intensity densities which in turn correlated positively with both mean peak flow cytometry intensities (Fig 3 i) and the molecular surface concentrations computed from fluorescence spectroscopy (Fig 3 j).

### 2.2 fluorescence-based inference of hybridization at doublet interfaces

In addition to demonstrating prescriptive PEGDPPE-based control over the molecular concentration of ssDNAPEGDPPE on cells prior to doublet formation, the agreement among the 3 fluorescence-based characterization methods allows us to make inferences about the molecular concentration of hybridizing molecules based on images of membrane profiles. To investigate the relationship between cohesion as we have computed it in our mechanical-geometric doublet model and the molecular surface concentration of ssDNAPEGDPPE at the cell-cell interface, we processed doublet images (n=50 per modification condition) utilizing the manually-measured cell center point data and radii to obtain automatically-isolated images of the cell-cell interfacial profile (Fig 4 a). These images were then converted to vertically aligned images. Observing the distribution of intensity along the length of these images, intensities were attenuated within 1 *μm* from the perimeter of the interface, but otherwise apparently uniform throughout the center region ie standard deviations less than 10% in pixel intensity for most cross-sections. The aligned images were converted to collapsed into 1D profiles (Fig 4 b) and integrated to obtain a single mass intensity value for each doublet interface. Across all tested PEGDPPE:FAMSeqAPEGDPPE ratios, we observed a positive correlation between individual doublet mass intensities and their corresponding *Ā*_*contact*_ (Fig 4 c). Importantly, when we analyzed cytochalsin D-treated doublets, we observed a similar positive correlation between mass intensities and their corresponding *Ā*_*contact*_ across the range of doublets analyzed, however with an apparently lower modulus than the non-cytochalsin D treated doublets.

We converted integrated intensity profiles to fluorescence intensity-based surface densities by dividing them by the interface diameter *a*_*ij*_ while assuming a constant confocal microscope sectioning depth. When the geometrically determined mean cohesion values for different PEGDPPE: ssDNAPEGDPPE ratios are plotted against intensity densities, a nonlinear trend emerges (Fig 4 d), and the plot of *Ā*_*contact*_ vs intensity density similarly reveals a nonlinear relationship. The intensity density of the cell-cell interface should correspond to the molecular concentration of fluorescein and thus to the concentration of oligonucleotides at the interface, however because doublet interfaces exhibited intensity densities spanning a range past the upper limit of those taken from images of single cell membrane profiles (Fig 4 e), we were unable to construct a reliable interpolative standard to infer the molecular concentration of FAMSeqAPEGDPPE at doublet interfaces and instead applied a less rigorous upward extrapolation (SFig 5). From Fig 4 e we can also see that the doublet intensity density curve is nonlinear compared to the linear shape of the single cell membrane intensity density curve.

## 3 Discussion and Conclusion

We used lipid-conjugated oligonucleotides to tune average intercellular cohesion in the absence of cadherins. A tomographic reconstruction of the 3D structure of a ssDNAPEGDPPE-cohered cell doublet revealed that fluorescently-labeled ssDNA was distributed throughout the interface between the two cells (Fig 1 b) in contrast, for example, to the annular distribution of cadherins at doublet interfaces observed by Engl et al^15^. This difference marks an aspect of ssDNAPEGDPPE-mediated adhesion which departs from adhesion modes that interface with the internal cytoskeleton and participate directly in an active force generating/sensing feedback loop. We examine ssDNAPEGDPPE doublets at a pseudo-steady state, though previous studies suggest changes would take place over longer timescales of hours such as the internalization of oligos^27^. The timescale of DNA hybridization is rapid, with the contact-formation phase occurring within seconds. Our system is also likely to involve actomyosin feedback in the form of cortex remodeling which has been shown to occur on timescales of minutes^42,43^ in response to cell shape change or external deformation. Our system therefore, while cadherin-free, is nevertheless not completely orthogonal to cytoskeletal dynamics.

Our model describes the space of geometric configurations (represented by *Ā*_*contact*_) in terms of Young’s modulus and cohesion, postulating effective independence (Fig 2 d). We hypothesized that modulation of one and/or the other could permit navigation over the configuration space. To explore this principle experimentally, we used a non-hybridizing PEGDPPE during ssDNAPEGDPPE modification to reduce the cohesion of cells and observed that the interfacial contact area decreased compared to doublets without PEGDPPE, and conversely cells treated with cortical actin-inhibiting cytochalasin-D exhibited large interfacial contact areas despite undergoing analogous ssDNAPEGDPPE modification as untreated cells (Fig 2 a, b, c). By preparing complementarily-modified cells with different PEGDPPE : ssDNAPEGDPPE ratios, we were able to systematically shift the distributions of interfacial contact areas determined from microscopy image analyses (Fig 2 e) and converting the measurements of doublet geometry into mean cohesion estimates via equation 6 show that systematic shifts in doublet geometry correspond to analogous shifts in the cohesion (Fig 2 f). For simplicity we assume a constant Young’s modulus independent of cortical rearrangement, conceiding that it is likely to be affected in the case of highly deformed doublets. Nevertheless we see that the PEGDPPE relative concentration enables prescriptive adjustment of cell-cell cohesion (Fig 2 g).

Our results apply to an ongoing discussion in cytoskeletal biology over what extent spontaneous separation of different cell types within tissues and the interfacial tension of multicellular domains are a result of differential adhesive compatibility analogous to phase separation of polar and nonpolar liquids^14,44–47^ relative to the cortical stiffness of cells^48,49^. A confounding factor comes from the annular actin reorganization in cadherin-cadherin-cohering cell doublets, which concentrates at the circumference of the cell-cell interface where cadherins also accumulate^15,50^. The reallocation of actin to the periphery of the contact reduces the compressive resistance that is normally characteristic of surface area minimizing spherical cortical tension. Without direct coupling to the internal cytoskeleton in our case, actin remodeling is likely restricted to that which occurs in response to compressive deformation. The treatment of doublets with cytochalsin-D under normal ssDNAPEGDPPE modification conditions and subsequent increased mean interfacial contact area demonstrates that the doublet geometry is influenced by the elastic mechanics of the cells. In a direct analogy to tissue surface tension, characterized by a minimization of the total exposed surface area multicellular aggregates to total aggregate volume ratio, the surface area to volume ratio of the cell doublet decreases with increasing interfacial contact area indicating a balance of cortical stiffness and cohesion in agreement with current models^51–53^.

In summary, we report on the establishment of a model to study cell-cell cohesion with a minimization of the active contribution of cohesion-inducing proteins like cadherins. By tuning the ratio of PEG-only and oligonucleotide-bearing lipid conjugates or by the introduction of stiffness-influencing pharmacological factors it was possible to affect the size of the contact area between cohering cell doublets. We use a mechanical model to describe the differences in contact area under different parameters. The system could be of value to studies which seek to examine the impact of cohesion without a major confounding influence of other cytoskeletal factors.

## 4 Acknowledgments

This study was supported by a Grant-in-Aid for Scientific Research on Innovative Areas “Nanomedicine Molecular Science” (No. 2306) and the Monbukagakusho Scholarship for graduate studies from the Ministry of Education, Culture, Sports, Science, and Technology (MEXT) of Japan.

