## supplemental_methods for "Tuning intercellular cohesion with membrane-anchored oligonucleotides"

### 1 Supplementary Data

#### 1.1 Numerical Generation of Theoretical of Steady State Interfacial Contact Area with Fixed Cell Volume Algorithm

The theoretical determination of interfacial contact areas for the steady state doublet model requires that the volume (highlighted in red in SFig 1 e) of each cell be conserved. With greater deformations due to greater cohesion, the radii of the intersecting spheres must change to compensate for the cap volume lost. Each cell volume $V_{celli}$ is defined by the difference of its sphere $V_i$ and its truncated cap $V_{capi}$.

$$V_{celli} = V_i - V_{capi}, V_{cellj} = V_j - V_{capj}. \tag{1}$$

The heights of the caps $h_i$ (shown in SFig 1 d) and $h_j$ can be represented in terms of the standard geometric parameters:

$$h_i = R_j - \sqrt{R_j^2 - \left(\frac{a_{ij}}{2}\right)^2}, h_j = R_i - \sqrt{R_i^2 - \left(\frac{a_{ij}}{2}\right)^2}. \tag{2}$$

The cap volumes are

$$V_{capi} = \frac{1}{3}\pi h_i^2(3R_i - hi), V_{capj} = \frac{1}{3}\pi h_j^2(3R_j - hj). \tag{3}$$

The cell volumes are then

$$V_{celli} = \frac{4}{3}\pi R_i^3 - \frac{1}{3}\pi h_i^2(3R_i - hi), V_{cellj} = \frac{4}{3}\pi R_j^3 - \frac{1}{3}\pi h_j^2(3R_j - hj). \tag{4}$$

Cell volumes are kept constant with numerical optimization as the interfacial areas are calculated according to the combined mechanical-cohesion model discussed in the main article. First, guess arrays of uniform radii $R_i$ and $R_j$ values equal to the radii corresponding to the volume of each cell in isolation, ie undeformed spheres. The length of the array is set to the length of the range of center-to-center distances $d_{ij}$ to be evaluated (for example from $\frac{R_i+R_j}{4}$ to $R_i + R_j$).

$$R_{guess} = \begin{pmatrix} R_{i_O} & R_{j_O} \\ R_{i_O} & R_{j_O} \\ \vdots & \vdots \\ R_{i_O} & R_{j_O} \end{pmatrix} \tag{5}$$

The force along the center-to-center axis is calculated according to Equation 5 from the main text as a function of the center-to-center distance $d_{ij}$ and for a given value of the cohesion $\gamma$ and the Young's modulus $\tilde{E}$. The root of this function is determined along the range of $d_{ij}$ values to determine the steady state geometry of the doublet, ie the $d_{ij}$ value of minimum interfacial energy and zero net force. The cell volumes at this point, according to Equation 1, will be underestimates of the initial cell volume as they are calculated from the uniform arrays in Equation 5. The sum squared error between the volume at the steady state point and the initial volume is minimized to readjust the values of Equation 5.

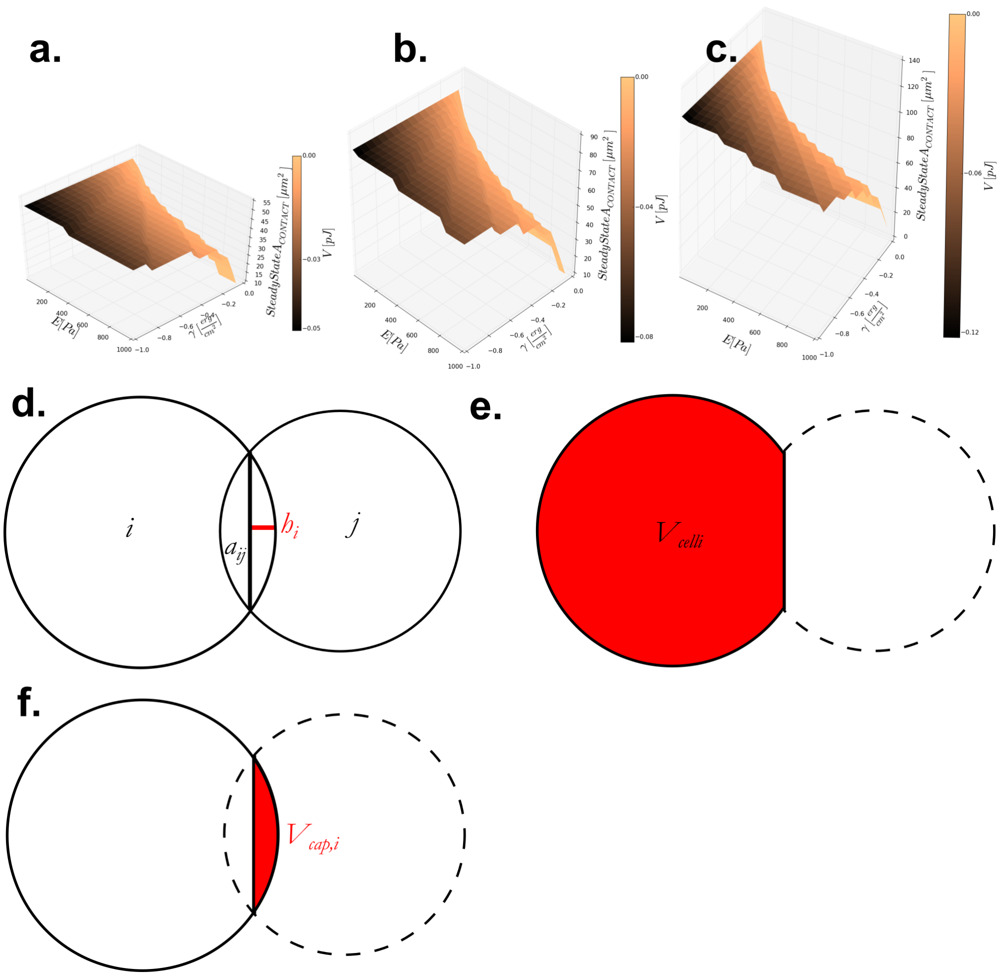


Figure 1: Volume conservation of cells during doublet mechanic mapping. (a) Contact area surface for cells of 4200 and 270 $\mu m^3$, (b) 3000 and 520 $\mu m^3$, (c) 2100 and 900 $\mu m^3$ (d) Geometric diagram of model doublet (e) Cell volume of a deformed sphere indicated in red (f) Truncated cap volume indicated in red.

The resulting array of radii vary with center-to-center distance while the volumes remain constant.

$$R_{adjusted} = \begin{pmatrix} R_i(d_{ij_0}) & R_j(d_{ij_0}) \\ R_i(d_{ij_1}) & R_j(d_{ij_1}) \\ \vdots & \vdots \\ R_i(d_{ij_n}) & R_j(d_{ij_n}) \end{pmatrix} \tag{6}$$

The process is repeated for a range of $\gamma$ and Young's modulus $\tilde{E}$ values to generate a map of steady state contact areas with the volumes of cells i and j kept constant as $V_{celli}$ and $V_{cellj}$. SFig 1 a-c shows the steady state contact area map for different cell volumes (4200 and 270, 3000 and 520, 2100 and 900 $\mu m^3$ respectively).

#### 1.2 Semi-automated doublet image processing to obtain geometric parameters and interfacial profiles

Confocal images of cell doublets (SFig 2 a and b) were processed first with a minimal manual algorithm consisting of drawing circles along the perimeter of the fluorescence cross sections. To identify perimeters reproducibly, fluorescence images were rendered in high dynamic range lookup tables. Although only one cell per doublet is labelled FAMSeqAPEGDPPE, weak autofluorescence (or diffused FAMSeqAPEGDPPE) permitted sufficient visibility to draw circles for both cells (SFig 2 c and d). From the two circles manually drawn, circle areas and central coordinates are then exported. Subsequent data processing is automated and proceeds independently of human judgement or bias.

From the areas and coordinates of manually drawn circles, $d_{ij}$, $R_i$, and $R_j$ are obtained through standard geometric relationships . From these three parameters, given Young's modulus and Poisson's ratio, and assuming a steady-state such that forces are balanced along the doublet axis, the mechanical-geometric model of the cell doublet is completely defined, and the cohesion can be calculated according to Materials and Methods: estimation of cohesion via doublet geometry.

Interfacial sections are then extracted automatically on the basis of the doublet geometry (SFig 2 e). The central point of the interfacial section is computed from one of the recorded coordinates of the circle centers (eg: $x_i, y_i$), the center-center axis angle $\theta_{CCA}$defined as the angle between the static x axis of the image and the line connecting cell centers, and the geometric parameters $d_{ij}$, $R_i$, and $R_j$:

$$x_c = x_i + \frac{(d_{ij}^2 - R_j^2 + R_i^2)}{2d_{ij}} cos\theta_{CCA}, \tag{7}$$

$$y_c = y_i + \frac{(d_{ij}^2 - R_j^2 + R_i^2)}{2d_{ij}} sin\theta_{CCA}. \tag{8}$$

A script in ImageJ uses coordinates computed from equations 7 and 8 for locating the central chord, defining a box around it with height truncated to the size of the chord length $a_{ij}$, and applying rotation by $\theta_{CCA}$ to extract interfacial profiles (SFig 2 f).

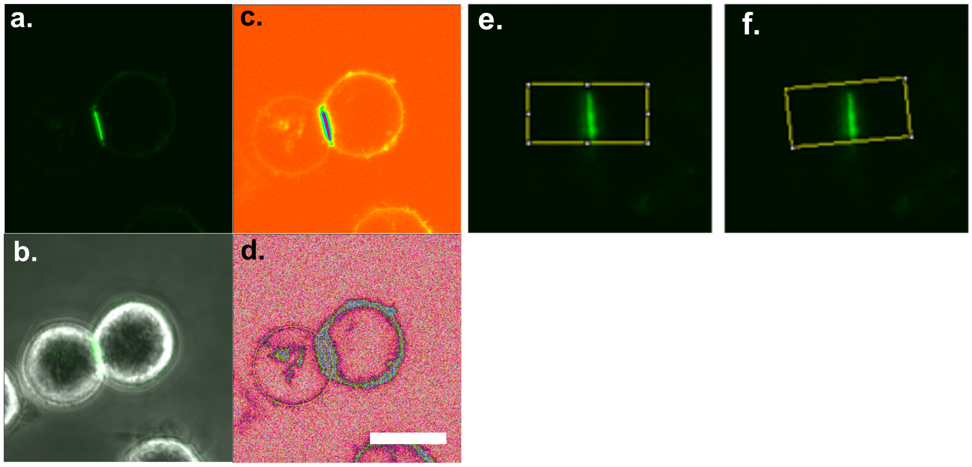


Figure 2: Extraction of interfacial profiles from confocal cross-sections of cell doublets. (a) Raw fluorescence image of a cell doublet with interface illuminated with FAMSeqAPEGDPPE. (b) Overlay of fluorescent image and phase contrast to show the existence of cohered cells joined by fluorescent interfacial region. (c) Fluorescent image from a. rendered in Spectrum color map for high dynamic range visualization. (d) Fluorescent image from a. rendered in Glasbey color map for ultra-high dynamic range visualization (Scale = 10 $\mu m$)(e) Box placement for image processing and recognition of the doublet interface. (f) Box angular adjustment based on the axis between the doublet centers.

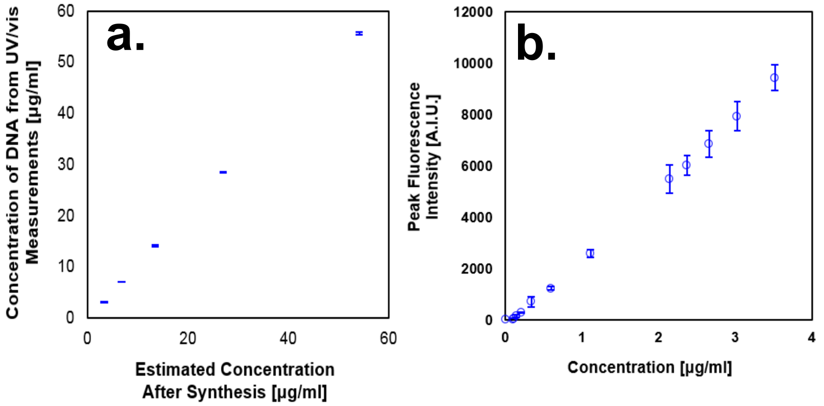


Figure 3: SeqA-PEG-DPPE standard.(a)UV/Vis concentration plotted versus synthesis concentration.(b) Peak fluorescence spectroscopy intensity plotted against Benzonase-treated FAM-seqA-PEG-DPPE.

#### 1.3 SeqA-PEG-DPPE Concentration Standard

In order to infer the concentration of FAM-SeqA-PEG-DPPE from the fluorescence emitted by liberated FAM, we calibrate concentration estimates according to the DNA content determined via absorbance at 260 $nm$. Absorbance was measured for samples of undigested FAM-SeqA-PEG-DPPE at 260 $nm$ and 495 $nm$ to compute spectra-corrected absorbance due to nucleotides:

$$A_{DNA} = A_{260} - fA_{495}, \tag{9}$$

where $A_{260}$ and $A_{495}$ are absorbances at 260 $nm$ and 495 $nm$ respectively, and $f = 0.279$ is the factor which corrects for the FAM contribution to absorbance at 260 $nm$.

The concentration of SeqA is related to $A_{DNA}$ by

$$c_{adjusted} = A_{DNA}(\mu_{SeqA}l)^{-1} = (A_{260} - fA_{495})(\mu_{SeqA}l)^{-1}, \tag{10}$$

where $\mu_{SeqA}$ and $l$ are the mass attenuation coefficient for SeqA and the cuvette path length respectively. The conversion from theoretical stoichiometric estimates of FAM-SeqA-PEG-DPPE concentration ($c_{stoichiometric}$) to adjusted concentration measurements related to DNA mass ($c_{adjusted}$) are obtained by linear regression of UV/Vis measurements at several concentrations (SFig 3 a) and is given by

$$c_{adjusted} = k_{correction}c_{stoichiometric} + \beta_{correction}, \tag{11}$$

where $k_{correction}$ and $\beta_{correction}$ are the constant of proportionality and intercept from linear regression respectively.

To relate fluorescence intensity measurements to concentration, a second standard curve was constructed by varying the concentration of FAM-SeqA-PEG-DPPE solutions treated with Benzonase to approach conditions relevant to cell experiments. In these experiments, triplicate spectra measurements were taken for multiple concentrations of FAM-SeqA-PEG-DPPE and plotted against adjusted concentration values (SFig 3 b). Linear regression yields

$$I_{520} = k_{spec}c_{supernatant} + \beta_{spec}, \tag{12}$$

where $k_{spec}$ and $\beta_{spec}$ are the constant of proportionality and intercept from linear regression respectively.

#### 1.4 Fluorescence spectroscopy of digestion-liberated FAM solutions from FAM-SeqA-PEG-DPPE-modified cells

Spectra of the various concentrations were measured at $n = 4$ separate binary dilutions ($DF$). Ensemble fluorescence values were calculated for a given sample of cells according to

$$I_{520} = \frac{1}{n}\sum_{DF=0}^{n=4} \bar{I}_{520}\left(\frac{1}{2}\right)^{-DF}, \tag{13}$$

where $\bar{I}_{520}$ is the average peak intensity over $i = 3$ replicate measurements.

Concentrations of SeqA molecules liberated in Benzonase extracts were calculated based on the linear FAM standard. Average molecular surface concentration on the cells $\bar{\rho}_{seqA}$ was estimated according to

$$\bar{\rho}_{seqA} = \frac{N_{seqA}}{n_{cells}\bar{A}_{cell}}, \tag{14}$$

where $\bar{A}_{cell}$ is the average cell surface area computed from spherical approximation with measured radii taken from the confocal cross sections at the point of a cell's maximum circumference, and $N_{seqA}$ is the measured number of seqA molecules calculated from concentration standard and known liquid volume of the extract. The number of cells $n_{cells}$ is estimated from the concentration of cells per unit volume measured by flow cytometry analysis.

#### 1.5 Membrane profile signal detection

To quantify membrane fluorescence intensity on a single-cell basis, images of confocal cross-sections of cells at their point of maximum circumference were analyzed. The centers of cells were identified by fitting circles to the membrane profile captured in each cross-section (Fig 4 a). A linear membrane profile was extracted by performing a polar to Cartesian transform relative to the fitted circle centers and 1 pixel corresponding to 1 angular degree (Fig 4 b). In order to reduce measurement error due to human judgement in the fitting of circles to membrane profiles, Cartesian profiles were processed such as to align the pixel of maximum intensity in each horizontal row along a single vertical axis (Fig 4 c). The aligned Cartesian profile intensities were then projected onto a 1D axis (Fig 4 d, e) and the peak signal was integrated and divided by 360 to yield a single quantity: fluorescence intensity per degree for each cell image. Images that underwent alignment had reduced deviation in integration value for a given displacement in circle center relative to those without alignment.

#### 1.6 Interface molecular concentration upward extraction from single cell profile data

Doublet interface intensities imaged in confocal cross-sections for a given meoPEGDPPE ratio were qualitatively greater than the corresponding single cell membrane intensities, thus the standard we attempted to produce for molecular surface concentration of SeqA based on differing single-cell membrane intensities can only be used as a rough estimate of doublet interface since upward extrapolation is required in cases where the observed interfacial intensity was greater than the upper limit observed with single cells (SFig 5 a). The theoretical hybridization surface energy can be computed from the extrapolated molecular concentration estimates combined with the nearest neighbor hybridization energy of SeqA. The cohesion estimated on the basis of doublet geometry and assumed elastic modulus scale positively with the total nearest neighbor hybridization energy per unit area (Fig 5 b).

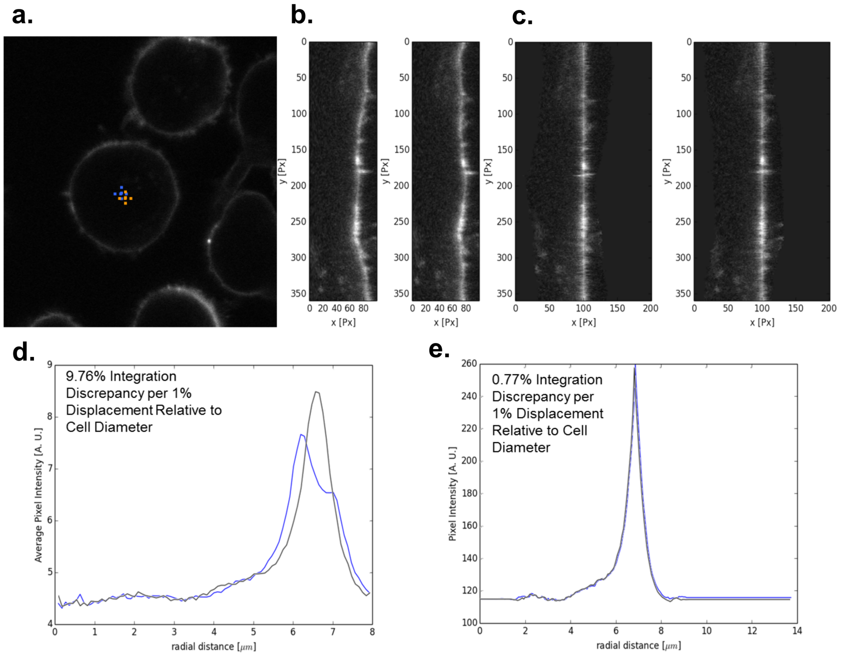


Figure 4: Image processing to extract intensity measurements from confocal images of single cells. (a) The centers of circles manually fitted to the membrane profile highlight in orange and blue. (b) Cartesian profiles corresponding to circles with the two centers from a.(c) Cartesian profiles from b with each row aligned according to its pixel of maximum intensity. (d) 1D projections of the unaligned profiles from b have a greater than 9 percent discrepancy in the integrated peak value for a 1 percent displacement in circle center. (e) 1D projection of the aligned profiles from c have reduced discrepancy relative to d.

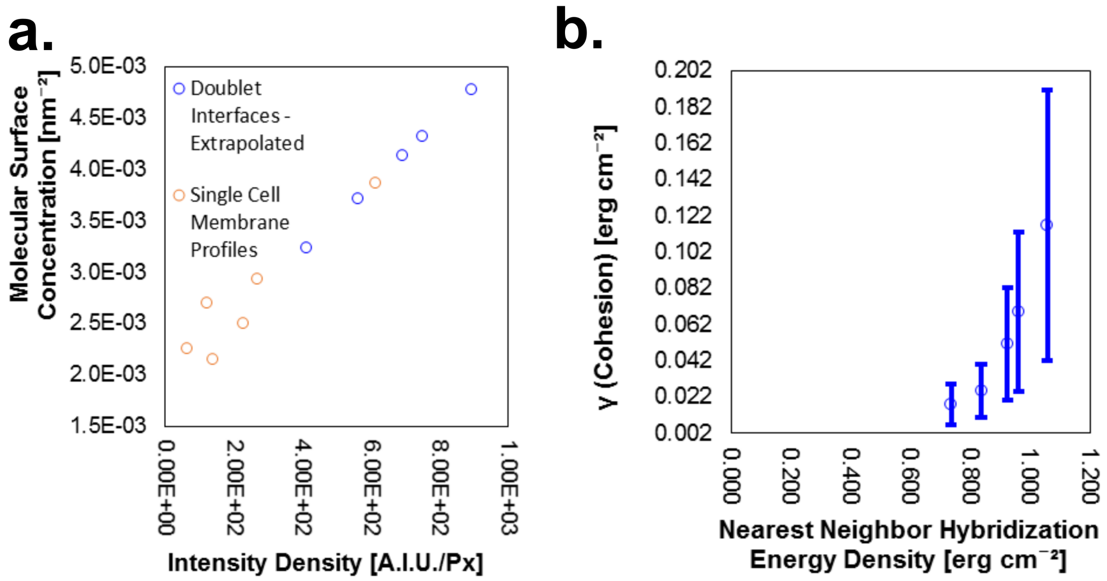


Figure 5: (a)Molecular surface concentration plotted versus intensity density measured from image analysis, with molecular surface concentration estimated for doublets on the basis of linear extrapolation from single cell measurements. (b) Model-derived cohesion plotted versus estimation of hybridization energy by nearest neighbor model of DNA hybridization.
